# Sexual dimorphism and plasticity in wing shape in three Diptera

**DOI:** 10.1101/135749

**Authors:** Micael Reis, Natalia Siomava, Ernst A. Wimmer, Nico Posnien

## Abstract

The ability of powered flight in insects facilitated their great evolutionary success allowing them to occupy various ecological niches. Beyond this primary task, wings are often involved in various premating behaviors, such as the generation of courtship songs and the initiation of mating in flight. These specific functions imply special adaptations of wing morphology, as well as sex-specific wing morphologies. Although wing morphology has been extensively studied in *Drosophila melanogaster*, a comprehensive understa nding of sexual wing shape dimorphisms and developmental plasticity is missing for other Diptera. Therefore, we raised flies of the three Diptera species *Drosophila melanogaster, Ceratitis capitata* and *Musca domestica* at different environmental conditions and applied geometric morphometrics to analyze wing shape. Our data showed extensive interspecific differences in wing shape, as well as a clear sexual wing shape dimorphism in all three species. We revealed an impact of different rearing temperatures wing shape in all three species, which was mostly explained by plasticity in wing size in *D. melanogaster*. Rearing densities had significant effects on allometric wing shape in *D. melanogaster*, while no obvious effects were observed for the other two species. Additionally, we do not find evidence for sex-specific response to different rearing conditions in all three species. We determined species-specific and common trends in shape alterations, and we hypothesize developmental and functional implications of our data.

**Contribution to the Field Statement:** The size and shape of organisms and organs must be tightly controlled during development to ensure proper functionality. However, morphological traits vary considerably in nature contributing to phenotypic diversity. Such variation can be the result of evolutionary adaptations as well as plasticity for example as reaction to changing environmental conditions during development. It is therefore a major aim in Biology to unravel the processes that control differences in adult morphology. Insect wings are excellent models to study how organ size and shape evolves because they facilitate basic tasks such as mating and feeding. Accordingly, a tremendous variety of wings sizes and shapes evolved in nature. Additionally, plasticity in wing morphology in response to different rearing conditions has been observed in many insects contributing to phenotypic diversity. In this work we applied Geometric Morphometrics to study wing shape in the three Diptera species: the Mediterranean fruit fly *Ceratitis capitata*, the Vinegar fly *Drosophila melanogaster* and the housefly *Musca domestica*. Flies were raised in different temperature and density regimes that allowed us to study the effects of these environmental factors on wing shape. Additionally, in accordance with different mating behaviors of these flies, we observed a clear sexual shape dimorphism in all three species. Since the three studied species represent serious pests and disease vectors, our findings may contribute to existing and future monitoring efforts.

## Introduction

Insects are the only group of arthropods that developed the ability of a powered flight. This adaptation allowed them to occupy various ecological niches, including air, and contributed to the morphological diversity and great ecological success of the entire class. Flying helps insects to surmount long distances in a relatively short time, facilitating basic tasks such as finding mating partners and food resources. In modern insects, wings became engaged in other essential processes such as mating and defense. For instance, numerous fruit flies use their wings to perform courtship songs, reflecting the size and vigor of males to help females choosing the right mating partner (Keiser et al., 1973; Burk and Webb, 1983; Webb et al., 1983b; Partridge et al., 1987). Some insects mate in the air during flight (Wilkinson and P. Johns, 2005), while others initiate the mating process in flight but always land prior to copulation (Murvosh et al., 1964). These different behaviors together with the intersexual food competition and various reproductive roles of the wing pose a constant selective pressure resulting in pronounced sexual dimorphism in wing size and shape in insects (Gidaszewski et al., 2009; Camargo et al., 2015). A thorough understanding of wing size and shape differences between sexes may thus allow linking environmental and behavioral constraints to morphological adaptations.

Variation in the size of distinct body parts, such as wings, across individuals of the same species is caused by differences in the genetic background, as well as by environmental cues (Beadle et al., 1938; Santos et al., 1994; Robinson and Partridge, 2001; Peck and Maddrell, 2005; DiAngelo et al., 2009; Shingleton et al., 2009; Siomava et al., 2016). Differences in developmental growth rates of an organ between sexes result in a sexual size dimorphism, which can be male- or female-biased if adult males or females are larger, respectively (Esperk et al., 2007; Fairbairn et al., 2007; Allen et al., 2011). The key component of the sexual size dimorphism is genetically defined in most insects (Blanckenhorn, 2000; Stillwell et al., 2010). However, in the light of developmental plasticity (i.e. the response of organ growth to different environmental conditions (Nettle and Bateson, 2015)), organ size may respond sex-specifically to different rearing conditions (Teder and Tammaru, 2005; Siomava et al., 2016). For instance, many exaggerated and sexually selected organs have been shown to grow disproportionally with body size in one sex in response to environmental cues (Bonduriansky, 2007; Lavine et al., 2015). Variation in organ size must be accompanied by shape changes to retain fully functional properties, suggesting that organ size and shape must be well-coordinated during development. Indeed, in *Drosophila melanogaster* wing size and shape are regulated by similar patterning and differentiation processes during larval and pupal wing imaginal disc development (Day and Lawrence, 2000; Matamoro-Vidal et al., 2015; Testa and Dworkin, 2016). Due to this tight morphological and developmental coupling, wing size and shape have been studied considered together for a long time (Cowley et al., 1986). Advances in geometric morphometrics methodology to quantify shape variation nowadays allow disentangling these two parameters and analyze size and shape independently (Bookstein, 1996; Bitner-Mathé and Klaczko, 1999; Debat et al., 2003; Mitteroecker and Gunz, 2009; Klingenberg, 2016). Such methods were successfully used to describe wing shape variation, as well as the impact of rearing conditions on wing shape in *Drosophila* and other insects (Cavicchi et al., 1991; Pezzoli et al., 1997; Moraes et al., 2004; Yeaman et al., 2010). Additionally, we have some understanding of sexual shape dimorphism in insect wings (Cowley et al., 1986; Gidaszewski et al., 2009; RIBAK et al., 2009; Allen et al., 2011; Benítez et al., 2011; Camargo et al., 2015; Gallesi et al., 2015; Virginio et al., 2015; Lorenz et al., 2017; Rodríguez and Liria, 2017). However, it remains largely unknown whether wing shape sex-specifically responds to different environmental cues and how such effects may be linked to wing size variation.

We have previously shown that the three dipteran species *Drosophila melanogaster, Ceratitis capitata*, and *Musca domestica* exhibit a clear sexual wing size dimorphism and that the response of wing size to different rearing conditions is sex dependent (Siomava et al., 2016). Therefore, these three species represent excellent models to test whether wing shape changes in a similar sex dependent manner. In this study, we applied geometric morphometrics to compare and comprehensively describe variation in wing morphology between *C. capitata, D. melanogaster* and *M. domestica*. We found clear evidence for sexual shape dimorphism in all three species with major contribution of wing size on shape differences in *D. melanogaster*. Different rearing temperatures had a strong effect on total and non-allometric wing shape in all three species, while density effects were most pronounced in *D. melanogaster*. Eventually, we did not find strong arguments for sexual dimorphism in response to the different rearing conditions, although *M. domestica* males showed slightly stronger differences than females. We identified highly variable regions, the radio-medial (r-m) crossvein, the R2+3 radial vein and the basal-medial-cubital (bm-cu) crossvein, which changed similarly among the three species in response to various larval densities and temperature regimes. This finding suggests that these regions may represent developmentally less robust wing compartments. We discuss our findings in the light of different mating behaviors in *C. capitata* and *D. melanogaster*, which produce courtship songs (Churchill-Stanland et al., 1986; Partridge et al., 1987), and *M. domestica*, which initiates mating in flight (Murvosh et al., 1964).

## Materials and Methods

### Fly species

All experiments were performed using three different fly species (Fig. 1A) (Siomava et al., 2016). We used the highly inbred laboratory strain *Drosophila melanogaster* w1118, which was kept at 18°C on standard food (400 g of malt extract, 400 g of corn flour, 50 g of soy flour, 110 g of sugar beet syrup, 51 g of agar, 90 g of yeast extract, 31.5 ml of propionic acid and 7.5 g of Nipagin dissolved in 40 ml of Ethanol, water up to 5 l). The other two flies were *Musca domestica* wild type ITA1 collected in Italy, Altavilla Silentia in 2013 (Y. Wu and L. Beukeboom, GELIFES, The Netherlands) and *Ceratitis capitata* wild type Egypt II (IAEA). The *M. domestica* strain was reared at room temperature (RT) (22±2°C) on food composed by 500 g of wheat bran, 75 g of wheat flour, 60 g of milk powder, 25 g of yeast extract, 872 ml of water and 18.85 ml of Nipagin (2.86 g of Nipagin in 10 ml of Ethanol). Adult *M. domestica* flies were provided with sugar water. *C. capitata* were kept at 28°C, 55 ± 5% RH on a diet composed by 52.5 g of yeast extract, 52.5 g of carrot powder, 2 g of Sodium benzoate, 1.75 g of agar, 2.25 ml of 32% HCl, 5 ml of Nipagin (2.86 g of Nipagin in 10 ml of Ethanol), water up to 500 ml for larvae. For adult flies, we used a 1:3 mixture of sugar and yeast extract.

**Fig. 1.**
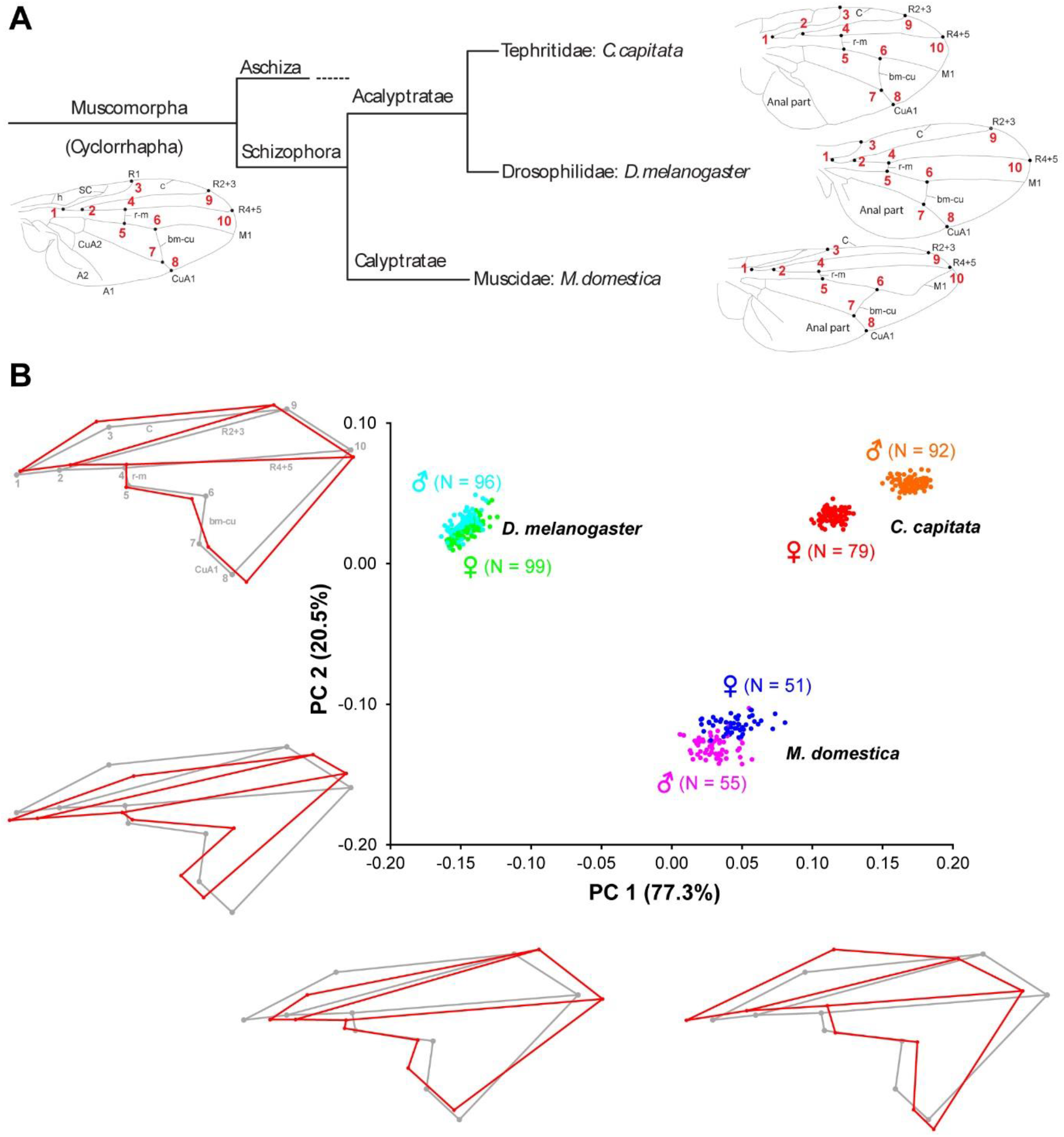
Schematic overview of the phylogenetic relationships Muscomorpha and interspecific wing shape variation *D. melanogaster, C. capitata and M. domestica*. (**A**) Homologous landmarks 1 to 10 in *C. capitata, D. melanogaster*, and *M. domestica* are shown as black points with the respective number in red. Vein abbreviations: A – anal vein; bm-cu – basal-madial-cubital crossvein; CuA – anterior cubital vein; C – costal vein; h – humeral crossvein; M – medial vein; R – radial veins; r-m – radio-medial crossvein; SC – subcosta. Branch lengths of the tree do not indicate evolutionary time or distance. (**B**) Wing shape variation between *D. melanogaster, C. capitata* and *M. domestica*. Principal Component Analysis of shape (PC1 and PC2). The red wireframes depict shape changes along the main axes of variation (PC1: −0.2, 0.2; PC2: −0.2, 0.1) relative to the grey wireframes (0 for both axes). The amount of variation explained by each PC is shown in brackets and the number of individuals analyzed for each sex is provided next to each species.

### Treatment of experimental groups

To generate a range of sizes for each species, we varied two environmental factors known to influence overall body size – temperature and density. Prior to the experiment, *D. melanogaster* flies were placed at 25°C for two days. On the third day, flies were moved from vials into egg-collection chambers and provided with apple-agar plates. After several hours, we started egg collection by removing apple-agar plates with laid eggs once per hour. Collected plates were kept at 25°C for 24 h to allow embryonic development to complete. Freshly hatched first-instarlarvae were transferred into 50 ml vials with 15 ml of fly food. Three vials containing 25 larvae each (low density) and three vials with 300 larvae each (high density) were moved to 18°C; the second set of six vials with the same densities was left at 25°C.

*C. capitata* flies were kept at 28°C and allowed to lay eggs through a net into water. Every hour, eggs were collected and transferred to the larval food. After 22 h, first-instar larvae were transferred into small Petri dishes (diameter 55 mm) with 15 ml of the larval food in three densities: 25 (low density), 100 (medium density) or 300 (high density) larvae per plate. Two plates of each density were moved to 18°C. The second set of six plates was left at 28°C for further development.

Eggs of *M. domestica* were collected in the wet larval food at RT and after 24 h, all hatched larvae were removed from food. Only larvae hatched within the next hour were transferred into 50 ml vials with 5 g of food. Collection of larvae was repeated several times to obtain two experimental sets with three replicates of three experimental densities 10 (low density), 20 (medium density) or 40 (high density) larvae. One set of nine vials was moved to 18°C, the other was left at RT.

After pupation, individuals of *C. capitata* and *M. domestica* were collected from the food and kept until eclosion in vials with a wet sponge, which was refreshed every other day.

The experimental temperature regimes were chosen for the following reasons. *D. melanogaster* is known to survive in the range 10 to 33°C, but flies remain fertile at 12 to 30°C with the optimum at 25°C (Hoffmann, 2010). Reproduction temperatures in *C. capitata* range from 14°C to 30°C with the optimum at 28°C (Duyck and Quilici, 2002; Navarro-Campos et al., 2011). *M. domestica* flies survive at 10 to 35°C (Hewitt, 1914) with the optimum between 24 and 27°C (Hafez, 1948; Chun-Hsung, 2012). The low temperature for our experiment was chose as the one above the survival and fertile minimums for all three species – 18°C. The warm temperature was aimed to be optimal for each species.

### Data collection

For each combination of temperature and density, we randomly picked at least four flies of each sex for *M. domestica* and 10 flies of each sex for *C. capitata* and *D. melanogaster*. Both wings were dissected, embedded in *Roti*®-Histokitt II (Roth, Buchenau) on a microscope slide, and photographed under the Leica MZ16 FA stereo microscope with the Q Imaging Micro Publisher 5.0 RTV Camera.

Wing shape was analyzed using landmark-based geometric morphometric methods (Rohlf, 1990; Bookstein, 1991). We digitized 10 anatomically homologous landmarks on wings of the three species (Fig. 1A). The landmarks were the following (nomenclature is given after (Colless and McAlpine, 1991)): 1, branching point of veins R_1_ and R_S_ (base of R_2+3_ and R_4+5_); 2, branching point of veins R_2+3_and R_4+5_; 3, intersection of veins C and R_1_; 4, intersection of vein R_4+5_ and crossvein *r-m* (anterior crossvein); 5, intersection of crossvein *r-m* and vein M_1+2_; 6, intersection of vein M_1+2_ and crossvein *i-m* (posterior crossvein); 7, intersection of crossvein *i-m* and vein M_3+4_; 8, intersection of M_3+4_ and the wing margin; 9, intersection of veins C and R_2+3_; 10, intersection of veins C and R_4+5_.

### Procrustes superimposition and interspecies comparison

Wing images were digitized using tpsUtil and tpsDig2 (Rohlf, 2015) to obtain raw *x* and *y* landmark coordinates. Using superimposition methods, it is possible to register landmarks of a sample to a common coordinate system in three steps: translating all landmark configurations to the same centroid, scaling all configurations to the same centroid size, and rotating all configurations until the summed squared distances between the landmarks and their corresponding sample average is a minimum scaling (Slice, 2005; Mitteroecker et al., 2013). To follow these three steps, we applied the generalized Procrustes analysis (GPA) (Dryden and Mardia, 1998; Slice, 2005) as implemented in MorphoJ (version 1.06d) (Klingenberg, 2011). The wings were aligned by principal axes, the mean configuration of landmarks was computed, and each wing was projected to a linear shape tangent space. The coordinates of the aligned wings were the Procrustes coordinates. To evaluate wing asymmetry, we used the R package Geomorph (v. 3.3.1) (Adams et al., 2021) to partition shape variation by individuals, side (directional asymmetry) and the interaction between individual and side (fluctuating asymmetry). Procrustes ANOVA was used to evaluate statistical significance. Since we obtained no evidence for fluctuating asymmetry for all species (Table S1), we averaged the coordinates for the right and left wings for further analyses. Next, we used Type III Procrustes ANOVA (with Randomization of null model residuals and 1,000 permutations) as implemented in Geomorph (v. 3.3.1) (Adams et al., 2021) to determine the effect of the species, sex, temperature, and density as well as potential interactions between these explanatory variables and shape (Table 1). Temperature and density values were re-labelled as low, high, or intermediate to allow the comparison between species. To visualize the main components of shape variation, we performed a Principal Component Analysis (PCA) and generated wireframes using MorphoJ (version 1.06d) (Fig. 1B).

**Table 1.** Effects of species, sex, temperature, density and their interactions on total wing shape tested by Type III Procrustes ANOVA.

|  | Df | SS | MS | R<sup>2</sup> | F | Z | P-value |
| --- | --- | --- | --- | --- | --- | --- | --- |
| spec | 2 | 0.844 | 0.422 | 0.081 | 1260.417 | 8.506 | **0.001** |
| sex | 1 | 0.027 | 0.027 | 0.003 | 80.926 | 7.175 | **0.001** |
| temp | 1 | 0.000 | 0.000 | 0.000 | 1.285 | 0.646 | 0.261 |
| dens | 2 | 0.001 | 0.000 | 0.000 | 1.399 | 0.957 | 0.168 |
| spec:sex | 2 | 0.036 | 0.018 | 0.003 | 54.078 | 9.158 | **0.001** |
| spec:temp | 2 | 0.004 | 0.002 | 0.000 | 5.356 | 4.178 | **0.001** |
| sex:temp | 1 | 0.000 | 0.000 | 0.000 | 0.897 | 0.106 | 0.471 |
| spec:dens | 2 | 0.003 | 0.001 | 0.000 | 3.701 | 3.253 | **0.001** |
| sex:dens | 2 | 0.001 | 0.000 | 0.000 | 1.313 | 0.815 | 0.205 |
| temp:dens | 2 | 0.001 | 0.000 | 0.000 | 1.061 | 0.266 | 0.408 |
| spec:sex:temp | 2 | 0.001 | 0.000 | 0.000 | 1.476 | 1.183 | 0.128 |
| spec:sex:dens | 2 | 0.002 | 0.001 | 0.000 | 2.551 | 2.387 | **0.005** |
| spec:temp:dens | 2 | 0.001 | 0.001 | 0.000 | 1.560 | 1.264 | 0.110 |
| sex:temp:dens | 2 | 0.001 | 0.000 | 0.000 | 0.753 | -0.530 | 0.700 |
| spec:sex:temp:dens | 2 | 0.001 | 0.001 | 0.000 | 1.537 | 1.245 | 0.107 |

Significant effects and interactions at $P \leq 0.01$ are provided as bold-italic P-values
spec – species, temp – temperature, dens – density, Df – degree of freedom, SS – Sum of Squares, MS – Mean Squares

### Intraspecific sexual dimorphism and effects of rearing conditions

Since most of the variation in shape was explained by differences between species (see Results), we split the analysis to further evaluate the effects of sex, rearing temperature, and density as well as potential interactions on wing shape within each species using Procrustes ANOVA in Geomorph (v. 3.3.1) (TableS2). Magnitudes of sexual shape dimorphism were estimated using the Discriminant Function Analysis (DFA) and expressed in units of Procrustes distance using MorphoJ (version 1.06d). DFA identifies shape features that differ the most between groups relative to within groups and it can only be applied to contrast two experimental groups. Therefore, we used this method to define sexual shape dimorphism (males and females), as well as effects of the rearing temperature (high and low) and density (high and low) in each species. Wing shape changes were visualized using wireframe graphs. To test for the significance of the observed differences, we ran a permutation test with 1,000 random permutations (Good, 1994) for each test using MorphoJ (version 1.06d).

### Estimation of the non-allometric component of shape

To evaluate the impact of wing size on shape variation, we estimated wing centroid size (WCS) that was computed from raw data of landmarks and measured as the square root of the sum of squared deviations of landmarks around their centroid (Bookstein, 1996; Dryden and Mardia, 1998; Slice, 2005) in MorphoJ (version 1.06d). The WCS values were corrected for differences in magnification and resolution among photos before being used as covariate. Since a Type III Procrustes ANOVA for all three species including WCS as covariate revealed significant interaction between species and WCS (TableS3, sheet “All Species_total”), we estimated the non-allometric shape component for each species independently. The non-allometric Procrustes coordinates were obtained as the residuals of the multiple regression of the Procrustes coordinates onto WCS pooling the samples by sex, temperature, and density, as implemented in MorphoJ (version 1.06d) (Klingenberg, 2011, 2016). To evaluate the success of the size correction, we ran a Type II ANOVA testing for additive effects of explanatory variables prior to and after the size correction. This analysis revealed that WCS had no impact on wing shape after size correction (TableS3, compare sheets “*_total” to “*_non-allometric”), suggesting that the approach accounted for most of the impact of size on shape. Note that species-specific Type III Procrustes ANOVAs including WCS as covariate revealed significant interactions for *C. capitata* and *D. melanogaster* (TableS3, sheets “C. capitata_total”, “D. melanogaster_total”), suggesting complex relationships between wing size and shape. Since the regression approach to estimate allometry across all explanatory variables (i.e. sex, temperature and density) assumes similar slopes of within-group regressions (Gidaszewski et al., 2009; Klingenberg, 2016), we analyzed those regressions in more detail. Although the directions of within-group regressions (i.e. regression scores of the regression of Procrustes coordinates onto WCS pooled by sex, temperature, and density) were comparable (Fig. S1), type III ANOVA testing for interactions of explanatory variables partially resulted in significant interactions in *C. capitata* and *D. melanogaster* (TableS4), suggesting some significant differences in slopes. Therefore, the distinction between allometric and non-allometric shape differences may not be fully accurate. However, due to a lack of a better method, we decided to proceed with the analysis and to interpret the results cautiously.

Magnitudes of non-allometric shape differences for each explanatory variable and for sex-specific effects of different temperature and density conditions were estimated using DFA as described before. Since wing size and shape of intermediate densities for *C. capitata* and *M. domestica* overlapped with the high- and low-density conditions, respectively (Fig. S2), we restricted the detailed shape analysis to the extreme values.

## Results

### Wing shape variation in dipteran species

To characterize wing shape variation among *Ceratitis capitata, Drosophila melanogaster*, and *Musca domestica*, we applied geometric morphometrics based on 10 landmarks (Fig. 1A). Procrustes ANOVA revealed that most of the shape variance was caused by differences among species (Table 1). This result was also reflected by the Principal Component Analysis (PCA) which clearly distinguished the three species with the first two Principal components (PCs) accounting for almost 98% of the variation (Fig. 1B). The main shape difference along PC1 (77.3% of the variation) reflected the ratio between the proximal and distal parts of the wing as well as wing length. The wireframe graphs showed that *C. capitata* wings were broad in the proximal part (landmarks 1-5) and narrow in the distal part (landmarks 6-10), while *D. melanogaster* wings showed the opposite pattern with the proximal part being heavily compressed along the anterior-posterior axis. *M. domestica* had an intermediate morphology being, however, more similar to *C. capitata* (Fig. 1B, wireframe graphs along PC1). PC2 explained 20.5% of the variation mainly accounting for the ratio between the length and width of the whole wing. *C. capitata* and *D. melanogaster* showed wider and shorter wings, while *M. domestica* showed more elongated wings (Fig. 1B, wireframe graphs along PC2). Together, we found evidence for extensive wing shape variation among *C. capitata, D. melanogaster*, and *M. domestica*.

### Sexual dimorphism in wing shape

Our interspecific shape analysis revealed a clear sexual shape dimorphism which was most pronounced for *C. capitata* (Fig. 1B, Table 1) (see also (Pretorius, 2005; Bogdanović et al., 2009; Camargo et al., 2015)). The extreme sexual shape dimorphism in *C. capitata* likely explains the significant interaction between species and sex (Table 1). Species-specific Procrustes ANOVA for all three species revealed that within-species shape variation was affected independently by sex and rearing conditions (i.e. temperature and density, see below) (TableS2). Therefore, we split the analysis by species and ran a Discriminant Function Analysis (DFA) to determine the total shape differences caused by sex (Fig. 2A-C). For all three species we found a highly significant sexual shape dimorphism (Fig. 2A-C). Male wings were broader than those of females in *C. capitata* (Fig. 2A) and *D. melanogaster* (Fig. 2B), while the opposite trend was observed in *M. domestica* (Fig. 2C). Wings of *C. capitata* females were slightly longer than male wings (Fig. 2A) and in *D. melanogaster* and *M. domestica* male wings were longer (Fig. 2B,C). The radio-medial crossvein (r-m) defined by landmarks 4 and 5 was different between males and females in *C. capitata* (Fig. 2A). In *D. melanogaster* we observed clear sexual differences in the radial vein R2+3 defined by landmarks 2 and 9 and in the basal-madial-cubital crossvein (bm-cu) defined by landmarks 6 and 7 (Fig. 2B). In summary, we found a clear total sexual shape dimorphism in all three studied species.

**Fig. 2.**
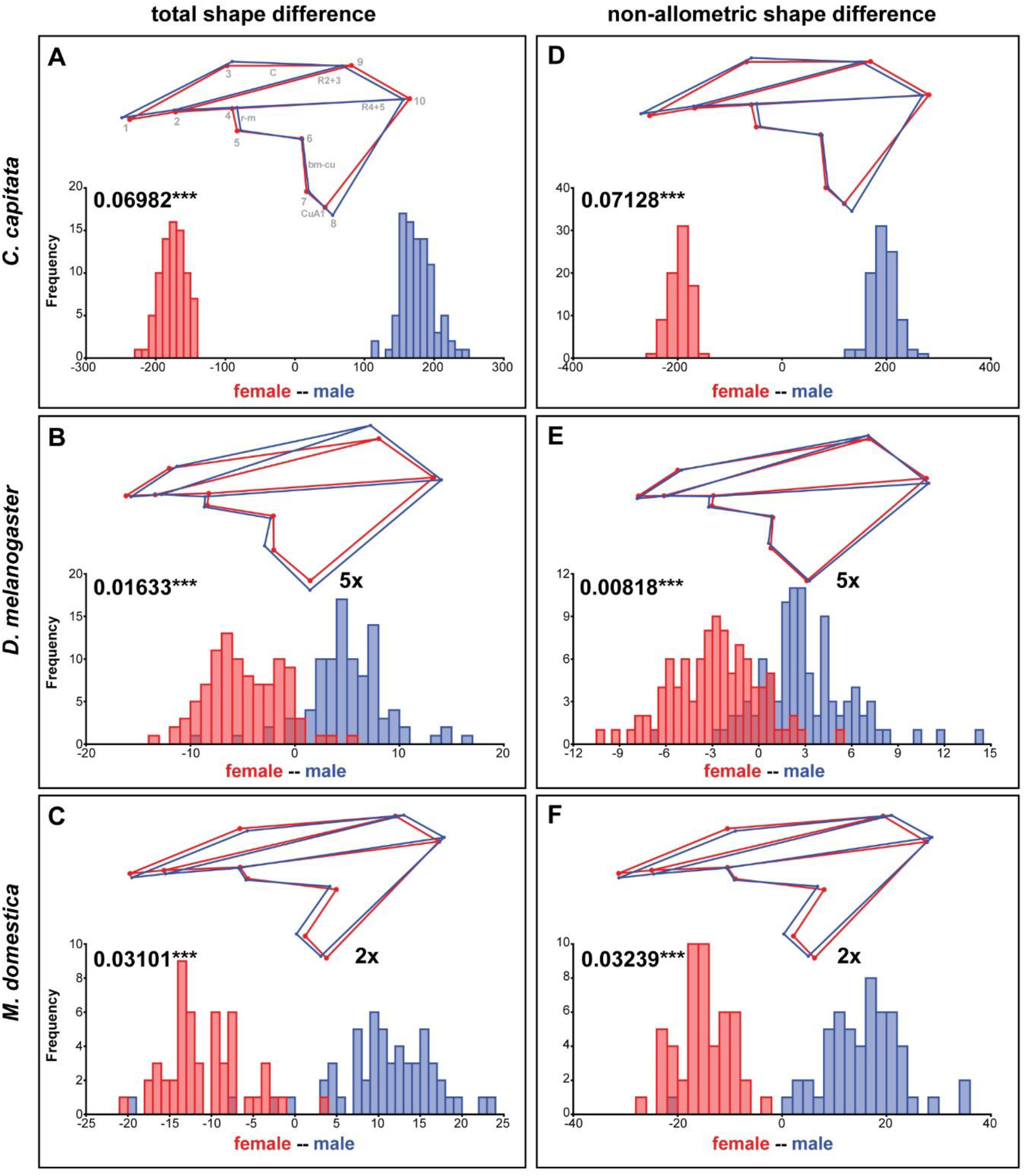
Total and non-allometric sexual shape dimorphism. Summary of Discriminant Function Analyses for total (**A-C**) and non-allometric (**D-F**) shape differences between females and males. The wireframes represent differences between female (red) and male (blue) average wing shapes. The scale factor is provided next to the wireframes. The magnitude of sexual shape dimorphism is indicated in units of Procrustes distance with the corresponding p-values based on 1,000 random permutations (***: P ≤ 0.001). Histograms with the distributions of the discriminant scores show shape separation into two distinct groups for each species. See also Table S5 for details.

### Influence of sexual size dimorphisms on wing shape

Since wings of the three studied species exhibit a clear sexual size dimorphism (Siomava et al., 2016) we next asked how much of the shape differences between sexes was explained by differences in wing size. Plots of the shape component that was most correlated with wing size against wing size implied a minor impact of size on wing shape in *C. capitata* and *M. domestica*, while wing shape was clearly associated with differences in wing size in *D. melanogaster* (Fig. S2). To analyze the non-allometric shape differences between sexes in more detail, we accounted for wing size (Klingenberg, 2016) (see Materials and Methods for details). A species-specific DFA using the non-allometric shape component revealed significant sexual shape dimorphism for all three species (Fig. 2D-F). This observation was confirmed by a Procrustes ANOVA (TableS2). The extant of the total compared to the non-allometric sexual shape dimorphism was very similar in *C. capitata* and *M. domestica* (Fig. 2D,F), suggesting that the allometric component had only a minor impact on wing shape in these two species. In *D. melanogaster* we found a clear impact of wing size on shape differences because the differences in shape decreased after size correction (Fig. 2E). Specifically, differences in the length and the width of the wings were largely reduced in the non-allometric shape component. These results suggest that major difference between sexes observed in *C. capitata* and *M. domestica* can be explained by the non-allometric shape component, while wing size differences contribute to the sexual shape dimorphism in *D. melanogaster*.

### Effect of different rearing conditions on wing shape

To evaluate the effect of wing size on shape in the three species in more detail, we raised flies at different temperatures and densities (Bitner-Mathé and Klaczko, 1999), which have been shown to influence body size. The Procrustes ANOVA for all three species revealed significant interactions between species and density and temperature, respectively (Table 1), suggesting species-specific effects of the rearing conditions on wing shape. Since temperature and density contributed additively to wing shape differences (TableS2), we therefore split the analyses by species and performed species-specific DFA to study the total shape differences caused by different rearing temperatures (Fig. 3A-C) and densities (Fig. 4A), respectively. Additionally, we calculated the non-allometric component of shape (see Materials and Methods for details) and analyzed the differences using DFA (Fig. 3D-F for temperature and Fig. 4B for density).

**Fig. 3.**
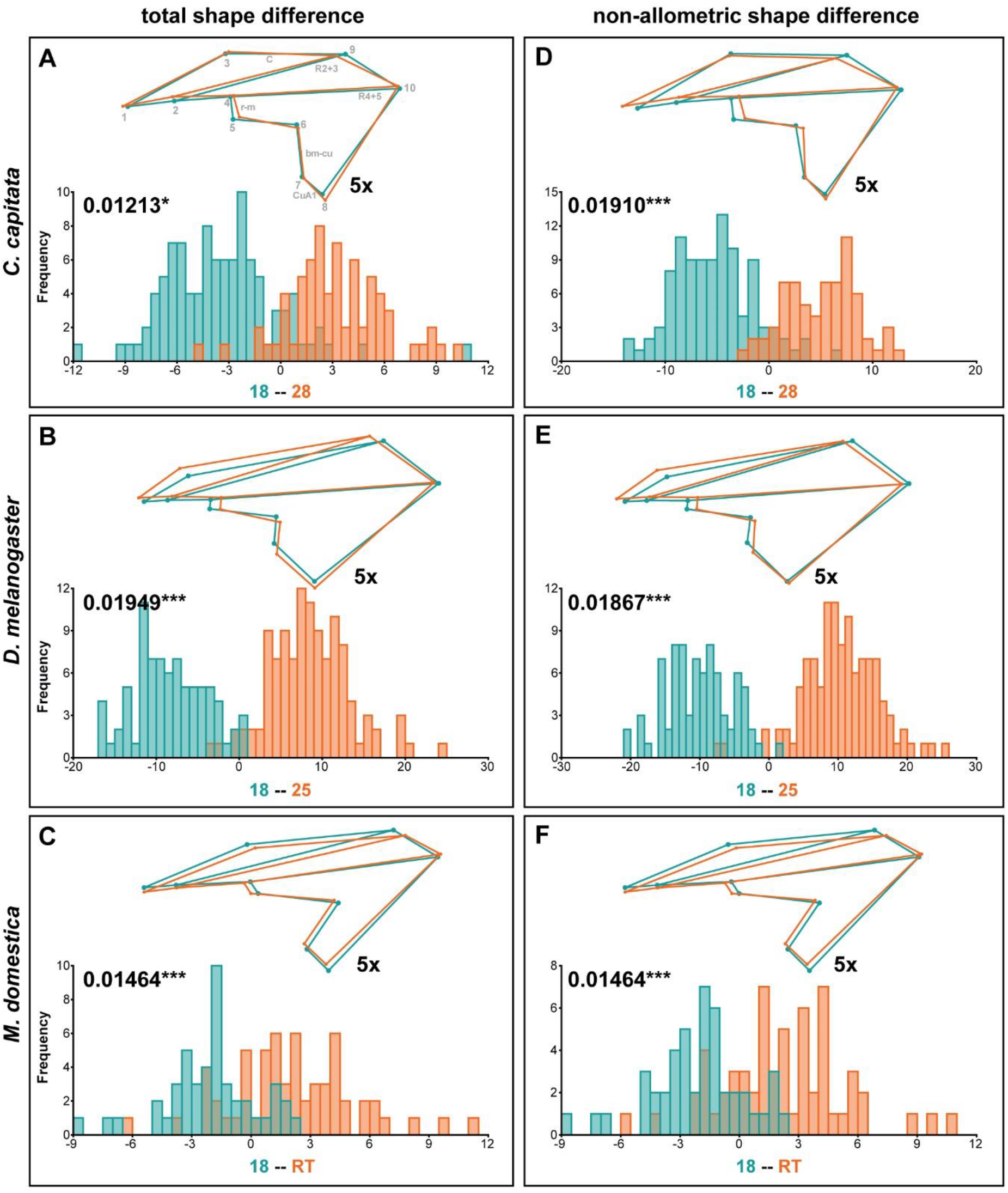
Total and non-allometric shape differences due to different rearing temperatures. Summary of Discriminant Function Analyses for total (**A-C**) and non-allometric (**D-F**) shape differences for flies raised at different temperatures. The wireframes represent differences between the low temperature (turquoise) and high temperature (orange) average wing shapes. The scale factor is provided next to the wireframes. The magnitude of shape variation is indicated in units of Procrustes distance with the corresponding p-values based on 1,000 random permutations (*: P ≤ 0.05; ***: P ≤ 0.001). Histograms with the distributions of the discriminant scores show shape separation into two distinct groups for each species. See also Table S5 for details.

**Fig. 4.**
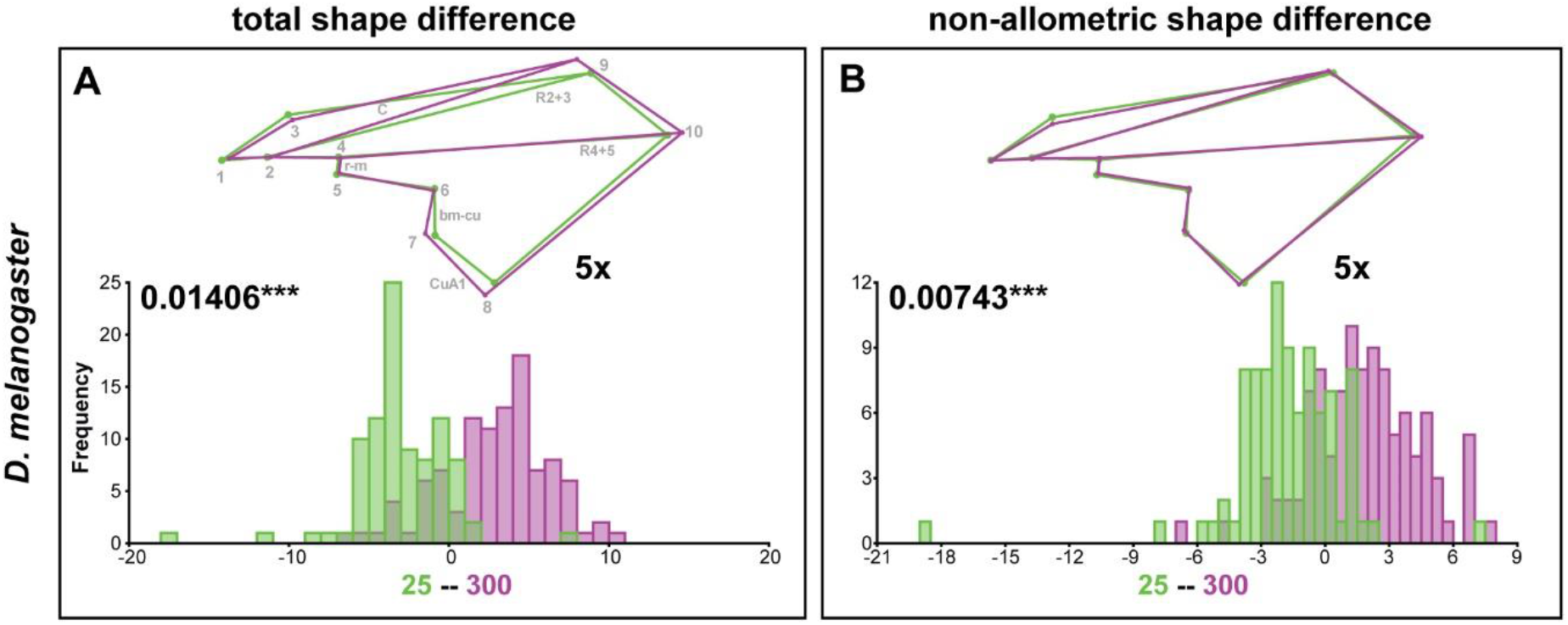
Total and non-allometric shape differences in *D. melanogaster* due to different rearing densities. Summary of Discriminant Function Analyses for total (**A**) and non-allometric (**B**) shape differences for flies raised at different densities. The wireframes represent differences between the low density (green) and high density (purple) average wing shapes. The scale factor is provided next to the wireframes. The magnitude of shape variation is indicated in units of Procrustes distance with the corresponding p-values based on 1,000 random permutations (***: P ≤ 0.001). Histograms with the distributions of the discriminant scores show shape separation into two distinct groups for each species. See also Table S5 for details.

In accordance with the species-specific Procrustes ANOVA testing for effects of rearing conditions on total wing shape variation (TableS2), the DFA clearly assigned wings to one of the two rearing temperatures (Fig. 3A-C). The most obvious effect of different rearing temperatures on wing shape was observed in *D. melanogaster* (Fig. 3B). Wings of *D. melanogaster* flies raised at higher temperatures were wider in the distal-central region defined by landmarks 7-9 and distally shortened (i.e. displacement of landmark 10) (Fig. 3B). The distal contraction remained after size correction, while the width was much less affected (Fig. 3E). This observation suggests that temperature-dependent plasticity in wing width is predominantly caused by differences in wing size. Different rearing temperatures also affected distal-centralwing width in *C. capitata* (Fig. 3A) and *M. domestica* (Fig. 3C). Narrower wings were observed at higher temperatures in *M. domestica*, while wings were narrower at lower temperatures in *C. capitata* (Fig. 3A,C). *M. domestica* wings were longer at higher rearing temperatures (i.e. displacement of landmark 10) (Fig. 3C). Only minor changes were present between total and non-allometric shape differences in *C. capitata* and *M. domestica* (Fig. 3D,F), suggesting that temperature-dependent size differences had only small effects on wing shape in these two species. In all three species, we observed temperature-dependent plasticity in the placement of the radio-medial (r-m) crossveins (defined by landmarks 4 and 5), the basal-medial-cubital (bm-cu) crossveins (defined by landmarks 6 and 7), the radial vein R2+3 (defined by landmarks 2 and 9) and the anterior cubital (CuA1) veins (defined by landmarks 7 and 8) (Fig. 3).

As suggested by the Procrustes ANOVA (TableS2), the DFA showed that different rearing densities only significantly affected total wing shape in *D. melanogaster* (Fig. 4A; Table S5). Wings of flies raised at high densities were wider in the central-distal region (defined by landmarks 7-9) and elongated (i.e. displaced landmark 10) (Fig. 4A). We observed plasticity in the placement of the basal-medial-cubital (bm-cu) crossveins (defined by landmarks 6 and 7), the anterior cubital (CuA1) veins (defined by landmarks 7 and 8), the radial vein R2+3 (defined by landmarks 2 and 9) and the costal (C) vein (defined by landmarks 3 and 9) (Fig. 4A). All these differences were gone after size correction (Fig. 4B), suggesting that wing shape differences in response to rearing densities were predominantly associated with variation in size. The displacement of landmark 3 was consistent in both analyses (compare Fig. 4A to 4B), implying that this region of the wing may be affected by different rearing densities independent of wing size. Despite a weak effect of density on *C. capitata* wing shape in our Procrustes ANOVA (TableS2), the DFA did not reveal significant shape differences for flies raised at high and low densities (Table S5). The same was true for *M. domestica*, supporting the Procrustes ANOVA results (TableS2). Note that we observed significant differences in wing shape at different rearing densities after size correction in *C. capitata* and *M. domestica* in our Procrustes ANOVA (TableS2) and in the DFA (Table S5).

In summary, we showed that different rearing temperatures had a profound effect on wing shape *D. melanogaster* and minor effects were observed in *C. capitata* and *M. domestica*. Variation in rearing densities only clearly affected wing shape in *D. melanogaster*.

### Sexual dimorphism in response to rearing conditions

Since the non-allometric component of shape was consistently affected by different rearing conditions (TableS2, Table S5), we next asked for each species, whether a sexual dimorphism in response to different rearing temperatures and densities exists. In all three species, we did not observe significant interactions between sex and rearing conditions in our Procrustes ANOVA of size corrected shape variables (TableS2). Accordingly, in *C. capitata* and *D. melanogaster* a DFA revealed very similar non-allometric shape differences between high and low temperatures (Fig. S3, Table S5) and densities (Fig. S4, Table S5), respectively for both sexes. Despite insignificant interaction terms in the Procrustes ANOVA the effect of interactions between sex and rearing conditions was up to 9 times higher in *M. domestica* compared to the other two species (see Sum of Squares in TableS2; “*_non-allometric”-sheets). Interestingly, the DFA revealed a significant effect of temperature (Fig. 5A,B, Table S5) and density (Fig. 5C,D, Table S5) on non-allometric shape differences in males, while females were unaffected.

**Fig. 5.**
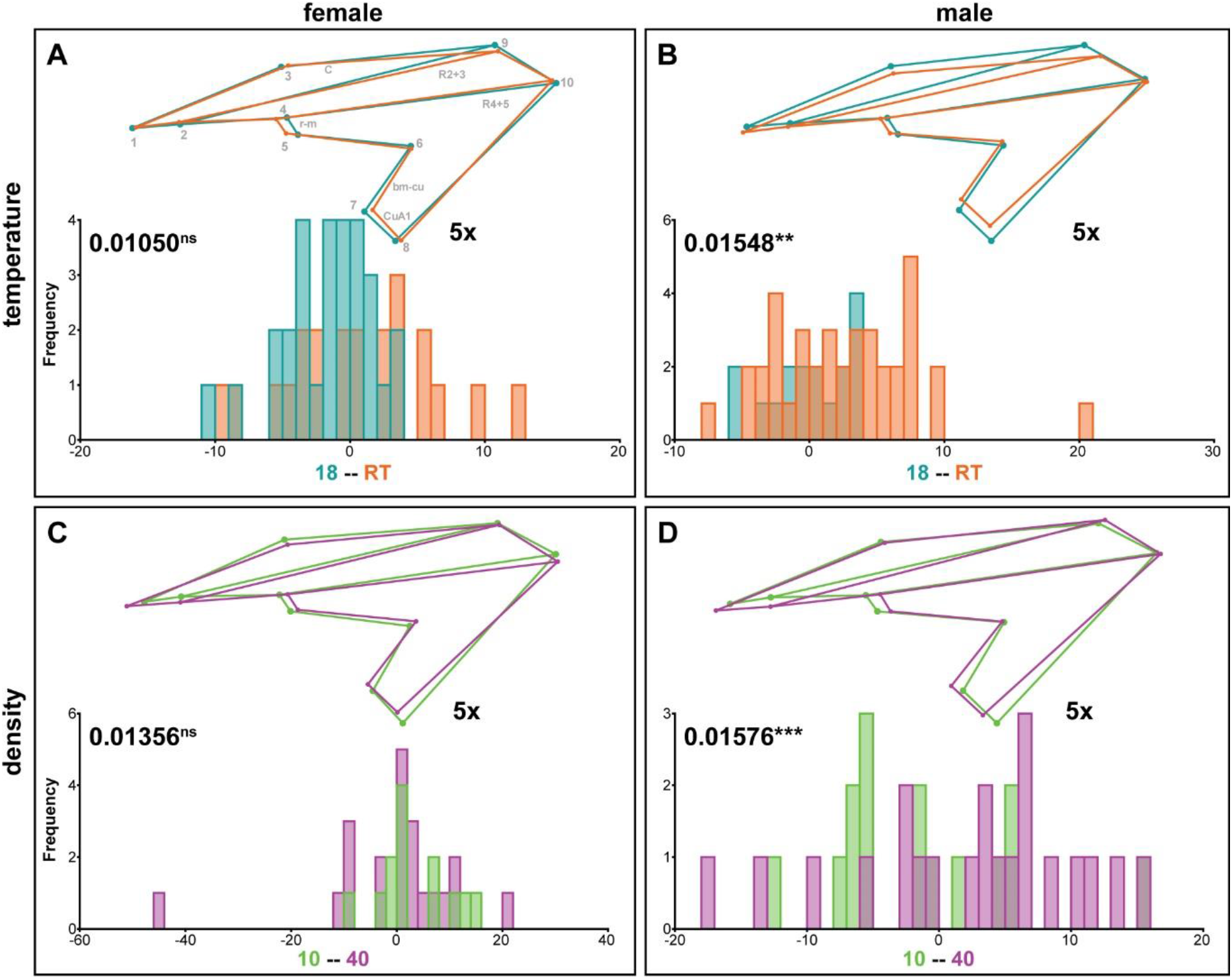
Sex-specific non-allometric shape differences due to different rearing temperatures and densities in *M. domestica*. Summary of Discriminant Function Analyses for non-allometric shape differences for flies raised at different temperatures (**A, B**) or densities (**C, D**). The wireframes represent differences between the low temperature (turquoise) and high temperature (orange) (**A, B**) and the low density (green) and high density (purple) (**C, D**) average wing shapes, respectively. The scale factor is provided next to the wireframes. The magnitude of shape variation is indicated in units of Procrustes distance with the corresponding p-values based on 1,000 random permutations (ns: P > 0.05; **: P ≤ 0.01; ***: P ≤ 0.001). Histograms with the distributions of the discriminant scores show shape separation into two distinct groups for each species. See also Table S5 for details.

In summary, we found no evidence for a sexual dimorphism in the response to different rearing conditions in *C. capitata* and *D. melanogaster*. In contrast, the non-allometric shape differences between extreme rearing temperatures and densities in *M. domestica* may be predominantly caused by affected males.

## Discussion

The comparison of wing shape among the three Diptera species *C. capitata, D. melanogaster* and *M. domestica* revealed extensive interspecific differences. The first two Principal Components (PCs) accounted for 97.8 % of the variation and captured differences in the wing width along the proximal-distal axis (PC1) and the ratio of wing width and wing length (PC2). In accordance with previous observations, we detected a clear sexual wing shape dimorphism in all three species (Bitner-Mathé and Klaczko, 1999; Lemic et al., 2020), which was most pronounced in *C. capitata*. Previous work on these three species revealed extensive sexual wing size dimorphism and it has been shown that males and females respond differently to different rearing conditions, which affect wing size (Siomava et al., 2016). Therefore, the major aim of this study was understanding the relationship between wing size and wing shape across sex and rearing conditions. In the following we will discuss our results with respect to size and shape relationships and we hypothesize developmental and functional implications of our findings.

### Total shape variation and size and shape relationships

Variation in organ size often entails shape changes (Debat et al., 2003; Klingenberg, 2016). Our analysis of the shape component that was most correlated with size variation showed a clear dependency on wing size in *D. melanogaster*, while the relationship was much less pronounced in the other two species (see data in Fig. S1 and Fig. S2). Based on our experimental design, the range of studied wing sizes was the result of a mix of effects of sex and rearing conditions (i.e. temperature and density) (Siomava et al., 2016). While the total number of analyzed flies per species was high, the number of flies for each subgroup (i.e. a certain sex at a specific rearing temperature and density, respectively) was rather low. Additionally, allometric relationships were different among subgroups (Fig. S1), suggesting that the regression-based approach to separate the allometric and non-allometric shape components may not be perfectly accurate (Gidaszewski et al., 2009; Klingenberg, 2016). However, despite these technical and analytical limitations, we could show that the three factors contributed additively to the observed shape variation and we confirmed that the size correction indeed removed the effect of centroid size on wing shape variation. Therefore, we are convinced that we could observe general trends, which will be briefly discussed separately for each factor.

### Sexual dimorphism

We observed a clear sexual shape dimorphism in all three species. While mostly wing width differed between females and males in *C. capitata* and *D. melanogaster*, the most obvious sexual differences in *M. domestica* was in wing length. According to a clear impact of sex on wing size (Siomava et al., 2016) (Fig. S2), we found a clear contribution of the allometric component to the shape difference between males and females in *D. melanogaster*. For instance, a shift of CuA1 along the wing margin as also described by (Bitner-Mathé and Klaczko, 1999), we could only detect when the allometric component was included. In general, exclusion of the allometric coefficient decreased the sexual shape dimorphism, suggesting that most of the observed shape differences could be explained by differences in wing size. In contrast, sex had only a minor effect on wing size in *C. capitata* and *M. domestica* (Siomava et al., 2016) (Fig. S2). Accordingly, the impact of the allometric component on wing shape was weak. For instance, the variation in the wing length that was explained by the allometric component in *D. melanogaster* was solely explained by the non-allometric component in *C. capitata* and *M. domestica*.

### Rearing temperature

Different rearing temperatures resulted in significant shape changes in all three species and the most obvious effects were observed on wing width, while wing length was only weakly affected. Wings of flies raised at higher temperatures were generally wider in *D. melanogaster* and *C. capitata* and narrower in *M. domestica*. In contrast to previous data that revealed a strong response of the distal wing region to different temperatures (Debat et al., 2003), we detected a slightly stronger plasticity in proximal landmarks (Fig. 3B, see e.g. landmarks 1 and 3) in *D. melanogaster*. This discrepancy may be due to the range of temperatures chosen in the different experiments. In the previous work, stressful conditions (12°C and 14°C as the cold temperature and up to 30°C as the high temperature) were included and wings of flies reared at extreme temperatures were clearly separated along CV2 in a Canonical Variate Analysis (CVA) (Fig. 3 in (Debat et al., 2003)). In contrast, we applied intermediate non-stressful regimes (18°C and 25°C), suggesting that the temperature effects on the distal wing region may partially be a stress response. It is important to note that we realized during our analysis, that room temperature (generally between 22 and 25°C) was likely lower than optimal for *M. domestica* (Siomava et al., 2016). Therefore, the obtained results for this species should be treated with some caution because we may not have covered entire range of optimal rearing temperatures and specifically for the Italian strain we used, the low temperature may elicit some stress response.

Different rearing temperatures had clear effects on wing size in *C. capitata* and *D. melanogaster* and no effect in *M. domestica* (Siomava et al., 2016) (Fig. S2). Accordingly, we observed differences between the non-allometric and the total shape differences across rearing temperatures in *C. capitata* and *D. melanogaster*, while wing shape plasticity was unaffected after size correction in *M. domestica*. For instance, we did not observe the displacements of the CuA1 and R2+3 veins after size correction, suggesting that these shape changes are associated with variation in wing size. Interestingly, our data suggests more pronounced shape differences after size correction in *C. capitata* (e.g. compare Fig. 3A to B). This finding is counterintuitive in the light of a partitioning of total shape variation into an allometric and a non-allometric component (Gidaszewski et al., 2009; Klingenberg, 2016). Given that the regression-based size correction applied in this work may not be optimal, the increased shape variation after accounting for wing size could be an artifact. However, a study in the *Leporinus cylindriformis* group of ray-finned fishes revealed that the distinction of species based on geometric morphometrics data was facilitated by the inclusion of size correction (SIDLAUSKAS et al., 2011), suggesting that indeed relevant aspects of shape variation could be recovered. In the future, more specific experiments with tightly controlled temperature regimes will help clarifying this aspect.

### Rearing density

Different rearing densities had only a significant effect on wing shape in *D. melanogaster*. Major effects were observed on wing width and to some degree also on wing length. Most of these differences were associated with density related plasticity in wing size (see also (Siomava et al., 2016); Fig. S2) because the shifts of the CuA1, bm-cu and R2+3 veins were not observable in the non-allometric shape component (compare Fig. 4A to B). Although we did not detect total wing shape differences in *C. capitata* and *M. domestica*, the non-allometric component of shape was significantly different among extreme rearing densities in both species. Whether this observation is an artefact of the size correction procedure remains to be established in future experiments.

### Sexual dimorphism in response to rearing conditions

Generally, we did not find evidence for a sexual dimorphism in the response to different rearing conditions in the three studied species. It is important to note that our experimental design is particularly limited with respect to this question because the number of individuals in each relevant group (i.e. split by sex, density and temperature) is very low and we might not have the statistical power to detect any general trends. However, our DFA may imply that non-allometric shape differences between extreme rearing temperatures and densities observed in *M. domestica* may be restricted to males. Future experiments addressing these specific questions should be performed to obtain more conclusive results.

### Developmental implications

Overall, we found a clear sexual dimorphism in wing shape in the three studied species. A tight connection of the shape variation and wing size differences in *D. melanogaster* suggests that the growth regulation, patterning, and differentiation processes (e.g. vein placement and axis determination) during the larval wing imaginal disc development may be also tightly linked. These processes, however, may be less connected and independently regulated in the other two species. A potential mechanism for the link between the increase in wing size and shape variation may come from the analysis of Ethiopian *D. melanogaster* populations that showed that increased wing size caused by high altitude was accompanied by a loss of buffering against environmental perturbations (Lack et al., 2016). It has recently been shown that the bone morphogenic protein (BMP) signaling pathways is buffered during embryonic dorsal-ventral axis formation by the action of the BMP-binding protein Crossveinless-2 and Jun N-terminal kinase (JNK) pathway activator Eiger (Egr) (Gavin-Smyth et al., 2013). Since the BMP signaling (Affolter and Basler, 2007; Wartlick et al., 2012; Akiyama and Gibson, 2015) and JNK pathway are crucial regulators of wing size in *D. melanogaster* (Willsey et al., 2016), these pathways are excellent starting points to study a potential connection between developmental buffering mechanisms and the concerted size and shape control during development.

In addition to changes in wing shape along the proximal-distal axis, our shape analysis revealed a high variation in the positioning of the r-m (landmarks 4 and 5), R2+3 (landmarks 2 and 9) and bm-cu (landmarks 6 and 7) veins that was common for all three species. Displacement of these veins was found in all temperature and density groups, suggesting that this region represents a very plastic aspect of wing patterning. Further support for this suggestion comes from the loss of buffering against environmental perturbations in high altitude Ethiopian *D. melanogaster* populations that showed an increase in wing size. It has been shown that this decanalization resulted in a higher level of abnormal wing development with various defects in crossvein development (i.e. incomplete, missing or additional crossveins) (Lack et al., 2016). Wing vein development requires a proper integration of various central signaling pathways, such as epidermal growth factor receptor (Egfr), Notch, bone morphogenetic protein (BMP), Hedgehog (Hh) and Wnt signaling (reviewed in Celis, 1998; Marcus, 2001; Blair, 2007). A potential link between these pathways and environmental perturbations has been suggested to be mediated by the heat shock protein Hsp90, since mutations in *Hsp83*, the gene coding for Hsp90, have been shown to result in various morphological abnormalities in adult traits, including the formation of additional crossveins (Rutherford and Lindquist, 1998). Furthermore, an in-depth analysis of the effect of varying levels of *Hsp83* expression on wing shape in *D. melanogaster* revealed that Hsp90 contributes to the buffering of developmental processes against environmental differences. Specifically, the placement of the r-m and bm-cu crossveins was affected to different degrees depending on the genetic background of the studied flies (Debat et al., 2006). Since crossveins develop later than longitudinal veins (Marcus, 2001), but still use similar signaling pathways (reviewed in Blair, 2007), these two vein types probably integrate the stress response differently either via Hsp90 or additional mechanisms. Additional factors are indeed likely because Debat et al. stated that Hsp90 alone cannot explain the entire canalization effect and they also propose the involvement of additional factors (Debat et al., 2006). More targeted experimental and molecular studies are necessary to address the developmental basis of the crossvein plasticity in more detail in insect species other than *D. melanogaster*.

### Potential functional implications of interspecific wing shape differences and intraspecific sexual wing shape dimorphism

Both size and shape of wings directly influence specific mating behaviors, such as flying or the generation of mating songs. In many dipteran fly species, male individuals produce species-specific courtship songs by fast and repetitive wing movements. For instance, Caribbean fruit fly females of *Anastrepha suspensa* judge the size and vigor of a potential mating partner by the intensity of its courtship song (Burk and Webb, 1983; Sivinski et al., 1984; Webb et al., 1984). Apparently, the intensity and audibility of these songs directly depend on the wing-beat frequency that the fly can afford for certain energy costs and wing fragility. In other fly species, such as *M. domestica*, the mating process is initiated during flight by an attack (“mating strike”) of a male against the back or side of a female. A successful “mating strike” usually results in the immediate landing and start of copulation (Murvosh et al., 1964).

Our morphometrics analysis revealed that *M. domestica* have wings that are longer and narrower than those of *C. capitata* and *D. melanogaster* (PC2) (Fig. 1B). This implies that *M. domestica* has the highest relative wingspan among the studied flies. The wing span is proportional to the moment of inertia (Shyy et al., 2013) and, thus, the moment of inertia should also the highest for *M. domestica* wings. Taking into account that the moment of inertia is inverse to the wing-beat frequency (Shyy et al., 2013), we conclude that the wing-beat frequency should be the lowest in *M. domestica* compared to the other two species. We also observed that male *M. domestica* wings were slightly elongated compared to female wings, additionally increasing the moment of inertia and required inertial power. Therefore, in accordance with the “mating strike” behavior in flight, *M. domestica* wings may be less suited for buzzing, but their wing shape might be under selection for better flight performance, that is facilitated by long and narrow wings (Alves and Bélo, 2002).

In contrast to *M. domestica, D. melanogaster* and *C. capitata* produce courtship songs by buzzing. Usually, their females favor males with a higher audibility (Churchill-Stanland et al., 1986; Partridge et al., 1987). Our study revealed that male wings were wider than female, shorter in case of *C. capitata*, and radial veins were more spread apart making wings more compact (Fig. 2). The allometric component of shape additionally increased this difference in *D. melanogaster*. The short and wide rounded wings of males in these species are likely to displace more air and repeat calling song pulses more quickly than long narrow wings, and the wing moment of inertia could be low enough to buzz. Interestingly, these flies produce two different types of wing vibration during the pre-mount courtship: the pulse song and sine song (Bennet-Clark et al., 1976; Rolli, 1976; Burk and Webb, 1983; Briceño et al., 1996; Briceño et al., 2002). The sine song is a continuous sinusoidal humming generated by small amplitude wing vibrations (Spieth, 1952; Ewing and Bennet-Clark, 1968; Hall, 1994; Markow and O’Grady, 2005; Ejima and Griffith, 2007). In *D. melanogaster*, its frequency ranges from 110 to 185 Hz (Wheeler et al., 1988), with the median value of approx. 160 Hz (Wheeler et al., 1988) or sometimes 130 Hz (Talyn and Dowse, 2004)^89^. In *C. capitata*, this frequency is similar to *D. melanogaster*, 165 Hz (Webb et al., 1983a). On the other hand, there is a difference in the pulse songs between these species. In case of *D. melanogaster*, it is composed of a series of single pulses (one to three cycles) separated by interpulse intervals. The frequency of these pulses is between 200– 280 Hz with the median value of 240 Hz (Cowling and Burnet, 1981; Rybak et al., 2002). Instead of pulses, *C. capitata* use a continuous vibration of wings when a male looks towards a female and keeps its abdomen bent ventrally (Briceño et al., 2002). An average frequency of such buzzing is about 350 Hz (Burk and Webb, 1983), which is almost half more frequent. Such high frequency of buzzing would require a lower moment of inertia, what could be achieved when the wing mass is concentrated near the axis of rotation (Berg and Rayner, 1995). Intriguingly, our shape analysis revealed major differences between *C. capitata* and *D. melanogaster* exactly in this region – variation in the width in proximal vs. distal regions (Fig. 1, PC1 axis). Thus, *C. capitata* had wider wings in the proximal part appropriate for high frequency buzzing and in *D. melanogaster* this part was narrower but perhaps wide enough for low frequency buzzing. Although this hypothesis remains to be tested, it is tempting to speculate that these shape differences may be linked to the mating behavior of the flies and specific properties of their courtship songs.

## Author Contributions

MR, NS, EAW and NP conceptualized the research; NS performed all fly experiments; MR and NS conducted data analysis and data visualization; NS, NP and MR wrote the original draft; all authors revised the manuscript; NP administered the project.

## Acknowledgments

We thank Y. Wu and L. Beukeboom for providing *M. domestica* flies. This work has been funded by a German Academic Exchange Service (DAAD) fellowship number 91540915 to NS, the Göttingen Graduate School for Neurosciences, Biophysics, and Molecular Biosciences (GGNB) and the Volkswagen Foundation (project number: 85 983; to NP).

## Data availability

Raw wing images, landmark files, R scripts and MorphoJ project files are available in an online repository (https://doi.org/10.25625/BXZHAF).

