## Supplementary Figures for "Sexual dimorphism and plasticity in wing shape in three Diptera"

##### This PDF file contains:

Supplementary Figures S1-S4

##### The following Supplementary Tables are available separately as Excel files:

**Table S1:** Results of Procrustes ANOVA testing for directional (Type II ANOVA) and fluctuating (Type III ANOVA) asymmetry.

**TableS2:** Species-specific Procrustes ANOVA testing for interaction and additive effects of variables for total shape and non-allometric shape components.

**TableS3:** Results of Procrustes ANOVA testing for additive effects of sex, rearing conditions and wing centroid size (Type II ANOVA), as well as their interactions (Type III ANOVA) on wing total and non-allometric wing shape, respectively.

**TableS4:** Species-specific Procrustes ANOVA testing for interaction of regression scores (allometries). Test for equality of slopes using the regression scores of the regression of Procrustes coordinates onto wing centroid size pooled by sex, temperature, and density. The regression scores correspond to the component of shape that is mostly correlated with variation in wing centroid size

**TableS5:** Summary of Discriminant Function Analyses (DFA) for pairwise comparisons of wing shape.

### Supplementary Figures

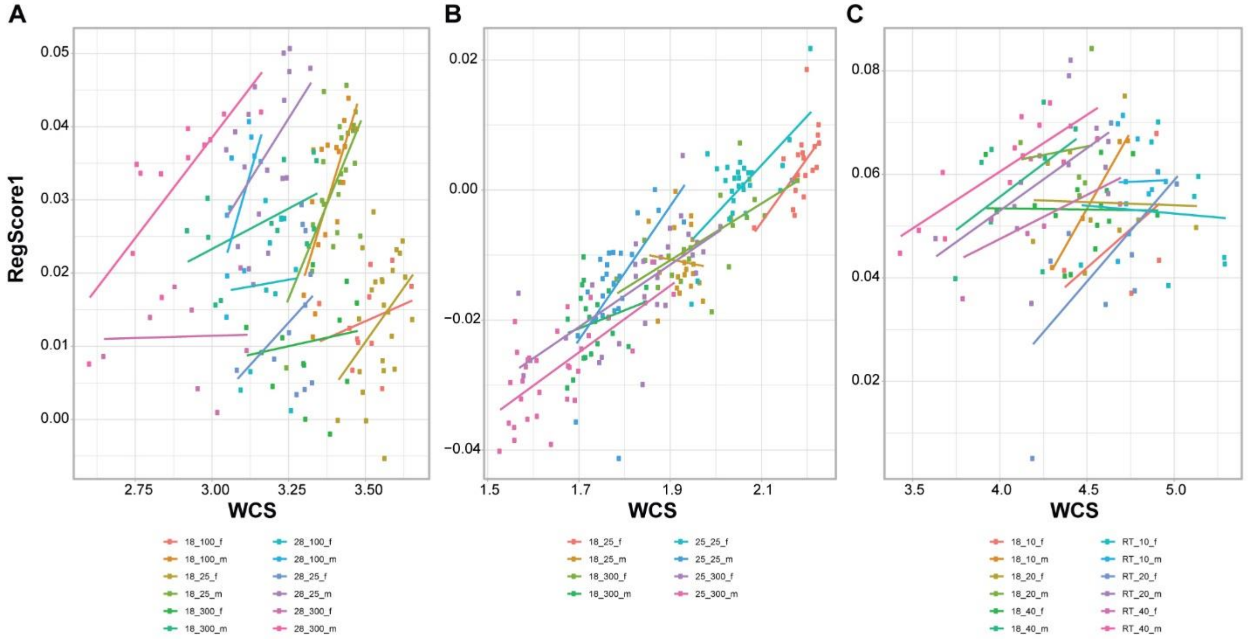


**Fig. S1. Relationship of the shape component that is most associated with wing size and wing centroid size for all categories.**
Scatterplots showing the relationship of the regression score (i.e. regression of Procrustes coordinates onto wing centroid size pooled by sex, temperature, and density) and wing centroid size (WCS). Linear regression lines were fitted for each category to illustrate the allometric relationships in detail. (**A**) *C. capitata*. (**B**) *D. melanogaster*. (**C**) *M. domestica*. The legend is defined as temperature_density_sex. f – females, m - male

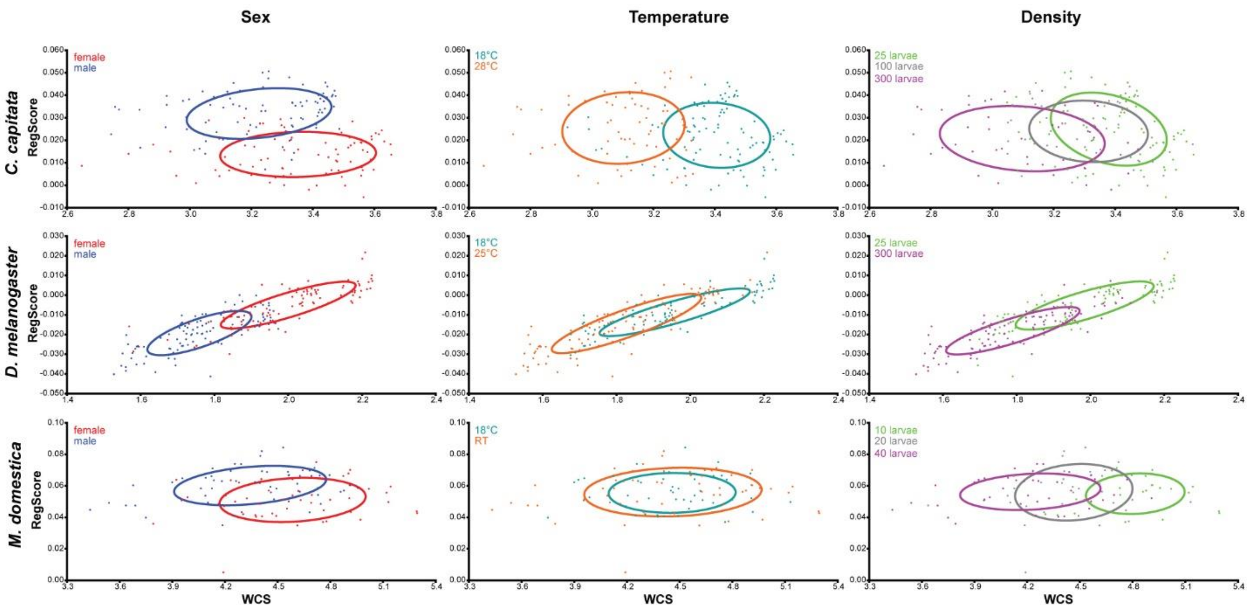


**Fig. S2. Relationship of the shape component that is most associated with wing size and wing centroid size pooled by sex, temperature and density.**
Scatterplots showing the relationship of the regression score (i.e. regression of Procrustes coordinates onto wing centroid size pooled by sex, temperature, and density) and wing centroid size (WCS). 90% confidence ellipses are shown for each group.

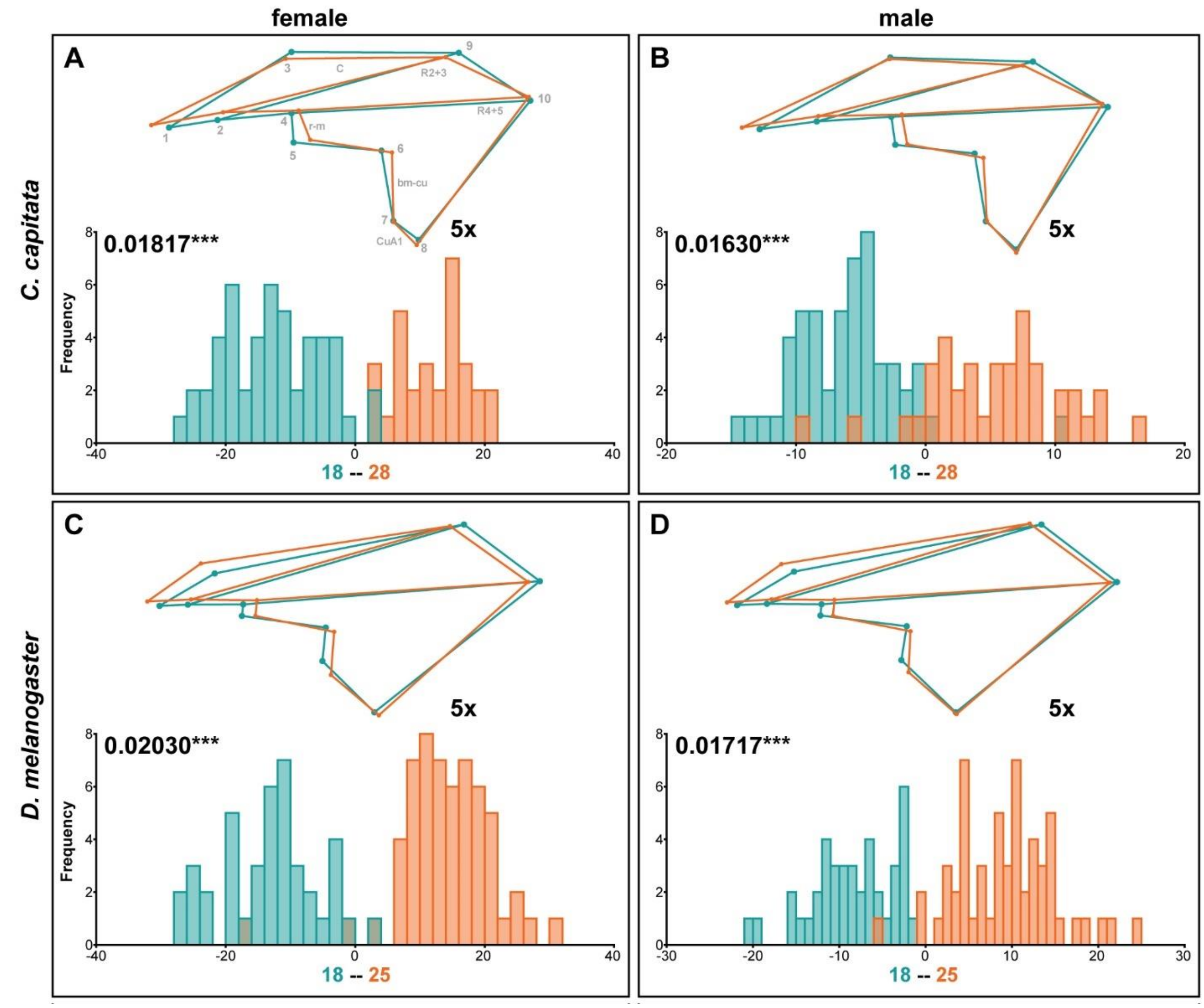


**Fig. S3. Sex-specific non-allometric shape differences due to different rearing temperatures**
Summary of Discriminant Function Analyses for non-allometric shape differences for flies raised at different temperatures for *C. capitata* (**A, B**) and *D. melanogaster* (**C, D**). The wireframes represent differences between the low temperature (turquoise) and high temperature (orange) average wing shapes. The scale factor is provided next to the wireframes. The magnitude of shape variation is indicated in units of Procrustes distance with the corresponding p-values based on 1,000 random permutations (***: $P \leq 0.001$). Histograms with the distributions of the discriminant scores show shape separation into two distinct groups for each species. See also Table S5 for details.

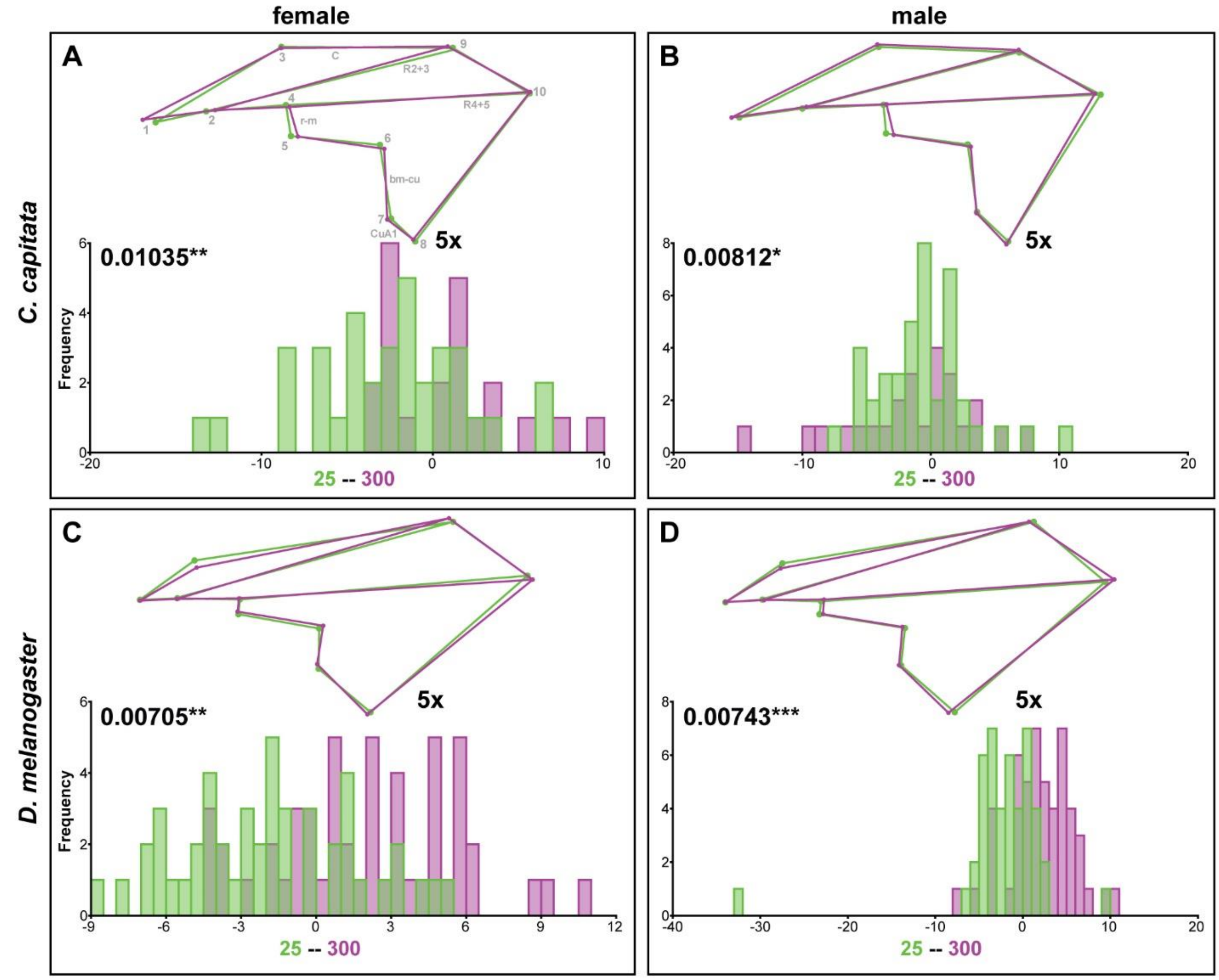


**Fig. S4. Sex-specific non-allometric shape differences due to different rearing densities**
Summary of Discriminant Function Analyses for non-allometric shape differences for flies raised at different densities for *C. capitata* (**A, B**) and *D. melanogaster* (**C, D**). The wireframes represent differences between the low density (grün) and high temperature (purple) average wing shapes. The scale factor is provided next to the wireframes. The magnitude of shape variation is indicated in units of Procrustes distance with the corresponding p-values based on 1,000 random permutations (*: $P \leq 0.05$; **: $P \leq 0.01$; ***: $P \leq 0.001$). Histograms with the distributions of the discriminant scores show shape separation into two distinct groups for each species. See also Table S5 for details.
